# Mitochondrial dysfunction and autophagy responses to skeletal muscle stress

**DOI:** 10.1101/597476

**Authors:** Anna S. Nichenko, W. Michael Southern, Anita E. Qualls, Alexandra B. Flemington, Grant H. Mercer, Amelia Yin, Hang Yin, Jarrod A. Call

**Affiliations:** Department of Kinesiology, University of Georgia, Athens, GA 30602, USA; Regenerative Bioscience Center, University of Georgia, Athens, GA 30602, USA; Center for Molecular Medicine, University of Georgia Athens, GA 30602, USA; Department of Biochemistry and Molecular Biology, University of Georgia, Athens, GA 30602, USA

## Abstract

Autophagy plays an important role in mitochondrial maintenance, yet many details of skeletal muscle autophagic activity are unresolved in the context of muscle stress and/or damage. Skeletal muscles from mice were stressed either by fatiguing contractions, eccentric contraction-induced injury (ECCI), or freeze injury (FI) to establish a timeline of mitochondrial function and autophagy induction after different forms of muscle stress. Only FI was sufficient to elicit a reduction in mitochondrial function (−88%, p=0.006), yet both ECCI and FI resulted in greater autophagy-related protein content (28-fold, p≤0.008) suggesting a tunable autophagic response. Muscles from another cohort of mice were used to determine specific forms of autophagy, i.e., flux and mitochondrial-specific, in response to muscle damage. Mitochondrial-specific autophagy was evident by accumulation of autophagy-related proteins in mitochondrial-enriched muscle fractions following FI (37-fold, p=0.017); however, autophagy flux, assessed by LC3II accumulation with the lysosomal inhibitor chloroquine, was insignificant suggesting a physiological bottleneck in the clearance of dysfunctional organelles following FI. Ulk1 muscle-specific knockout (Ulk1 MKO) mice were used to determine if autophagy is necessary for the recovery of mitochondrial function after muscle damage. Ulk1 MKO mice were weaker (−12%, p=0.012) and demonstrated altered satellite cell dynamics (e.g., proliferation) during muscle regeneration after FI compared to littermate control mice, but determination of autophagy necessity for the recovery of mitochondrial function was inconclusive. This study concludes that autophagy is a tunable cellular response to muscle damaging stress and may influence muscle fiber regeneration through interaction with satellite cells.

**Key Points Summary:**

- Muscle contractility dysfunction is well characterized after many different types of muscle stress however, the timing and magnitude of mitochondrial dysfunction and autophagy induction after different types of muscle stress is largely unknown.
- In this study we found that only traumatic freeze injury causes mitochondria dysfunction compared to fatigue contractions and eccentric contraction-induced injury, and that the autophagic response to muscle stress scales to the magnitude of muscle damage, i.e., freeze vs. eccentric contraction-induced injury.
- We determined that total autophagy-related protein content has a greater response to muscle fiber damage compared to autophagy flux likely reflecting a bottleneck of autophagosomes awaiting degradation following muscle injury.
- Using a skeletal gmuscle-specific autophagy knockout mouse (Ulk1), we found that muscle contractility and satellite cell activity might be influenced by cellular events within the adult muscle fiber following muscle damage.

## Introduction

The time course of muscle contractility loss after multiple types of muscle injury has been well documented (6, 22, 27, 39, 40), but the timing and severity of mitochondrial dysfunction after injury is largely unknown. Elucidating the loss and recovery of mitochondrial function is important because mitochondria provide crucial energy for satellite cell proliferation and differentiation, for the remodeling of damaged muscle fibers, and for the repair of initial membrane disruption (13, 37). Mitochondria are affected by muscle fiber damage, as we and others have reported a decrease in mitochondrial content and a subsequent rise in mitochondrial biogenesis during muscle regeneration (6, 9, 27, 36). However, the approximation of mitochondrial function using markers of mitochondrial content and biogenesis are inadequate to characterize mitochondrial function particularly when evaluating pathological conditions (23). Mitochondrial function is most appropriately analyzed by assessing the organelle’s ability to consume oxygen (i.e., mitochondrial respiratory function), and a primary goal of this study was to investigate the time course of mitochondrial dysfunction and recovery after various forms of muscle fiber stress.

Traditionally, the most physiologically-relevant marker of muscle fiber stress is a temporary or prolonged loss of contractility. There are many different types of muscle stressors that induce a temporary or prolonged loss of contractility (i.e. fatigue, eccentric contraction-induced injury, contusion, freeze injury, myotoxic injury, burn injury, and volumetric muscle loss injury) and, consequently, the severity and mechanism of reduced muscle contractility is unique for each stressor (39). Severe stressors, such as freeze or myotoxic injuries, destroy the contacted muscle fibers predominately by damaging the sarcolemma and disrupting intramuscular ion homeostasis which leads to a 65%-80% loss of contractility (14, 21, 39). Mitochondrial content follows a similar decline after these severe stresses (6, 27, 36), but the functional deficit (i.e., mitochondrial respiration) has not been adequately investigated. Less severe muscle stressors, like eccentric contraction-induced injuries, mainly disrupt excitation-contraction coupling and result in an initial 40%-60% decline in muscle contractility (38). There have been conflicting reports of mitochondrial oxygen consumption rates after downhill treadmill running, a mild and indirect form of eccentric contraction-induced injury in mice with some reporting no changes in mitochondrial respiration and others reporting transient changes immediately and up to 48 hours after the injury (24, 30, 31, 33). Additionally, this injury model is reported to elicit oxidative damage in the form of a greater presence of protein carbonyls and oxidized lipids that could implicate mitochondrial dysfunction (26, 33). Finally, even a mild muscle stressor such as muscle fatigue has been suggested to cause mitochondrial damage. Laker & Drake et al. recently published that horizontal treadmill running was associated with greater oxidation of the *pMitoTimer* reporter gene tagged to the Tyr-65 residue of Cytochrome C Oxidase subunit VIII (20), indicative of mitochondrial oxidative damage. Therefore, it is apparent that multiple types of muscle stressors may induce unique mitochondrial responses, although how relevant these are to mitochondrial function after muscle stress is unknown.

We have previously highlighted that when mitochondria are stressed or damaged a primary cellular mechanism for maintaining the quality of the mitochondria network is macroautophagy (6, 27). Macroautophagy (hereafter referred to as autophagy) is a cellular process which degrades dysfunctional organelles and proteins into their original amino acid and fatty acid components to be recycled in the cell. The Ulk1 (Unc-51 like autophagy activating kinase 1) complex initiates autophagy by signaling the Beclin1 (Atg6) complex to convert microtubule-associated protein light chain B I (LC3I) into LC3II, which then forms a double-membrane vesicle called an autophagosome. Autophagosomes encapsulate damaged organelles or proteins and eventually fuse with a lysosome to undergo degradation. Conventionally, measurements of Beclin1 or LC3 protein contents have been used to characterize broad changes in autophagy after various muscle stressors, but autophagy is a dynamic process that can be affected at many different stages, therefore the static measurements of Beclin1 and LC3 fail to capture the changes in overall autophagic flux (i.e., ongoing autophagic degradation) (17). Furthermore, the specific degradation of mitochondria by autophagy, alternatively defined as mitophagy, has primarily been investigated through localization of mitochondrial markers and LC3 puncta in skeletal muscle after treadmill running (20). There is currently a knowledge gap in the literature regarding the extent to which muscle fiber damage results in greater autophagy flux, and whether autophagy contributes to elimination of damaged mitochondria. Understanding the specific role of autophagy after muscle fiber damage may be leveraged to develop targeted therapeutic modalities to address muscle regeneration in conditions such as aging and muscular dystrophy where deficits in muscle repair and autophagy have been reported (25, 29, 32).

The objectives of this study were: 1) to elucidate the relationship between mitochondrial dysfunction and autophagy induction after different types of muscle stress; 2) to determine the extent to which autophagy flux and mitochondrial specific autophagy respond to muscle fiber stress; and 3) to determine if Ulk1-mediated autophagy is necessary for the recovery of contractile and mitochondrial function after muscle damage. We hypothesized that autophagy responses would scale to the magnitude of muscle stress, that autophagy flux and mitochondrial-specific autophagy would contribute to the autophagy response to muscle stress, and that Ulk1-mediated autophagy would be necessary for the recovery of muscle contractility and mitochondrial function after muscle stress.

## Methods

### Ethical Approval

All animal protocols were approved by the University of Georgia Animal Care and Use Committee under the national guidelines set by the Association for Assessment and Accreditation of Laboratory Animal Care.

### Animal Models

Male and female C57BL/6J mice aged 3-4 months were bred in-house and housed 5 per cage in a temperature-controlled facility with a 12:12 hour light:dark cycle. Muscle-specific Ulk1 knockout mice (Ulk1 MKO) with *myogenin-Cre* and LoxP flanked *Ulk1*and their *myogenin-Cre* negative littermates (LM) were used to test the necessity of Ulk1 for mitochondrial function and strength recovery after traumatic freeze injury. All mice had *ab libitum* access to food and water throughout the experiments.

### Experimental Design

The first cohort of wildtype C57BL/6J mice were used to assess the time course of mitochondrial function and autophagy induction after various muscle stressors. Briefly, mice were randomized into 3 groups; (i) a non-damaging metabolically fatiguing challenge (n=12), (ii) eccentric contraction-induced injury (n=20), and (iii) traumatic freeze injury (n=28). Tissues were analyzed for mitochondrial function, mitochondrial content, and autophagy induction immediately, 6 hours, and one day post for all groups. No additional time points were assessed for the fatigue group because initial data indicated there were no significant changes (Fig. 1 & 2). Additional time points, 3 and 7 days post, were assessed for the eccentric-contraction induced and freeze injury groups to characterize changes in mitochondrial function during the first week of recovery.

**Figure 1.**
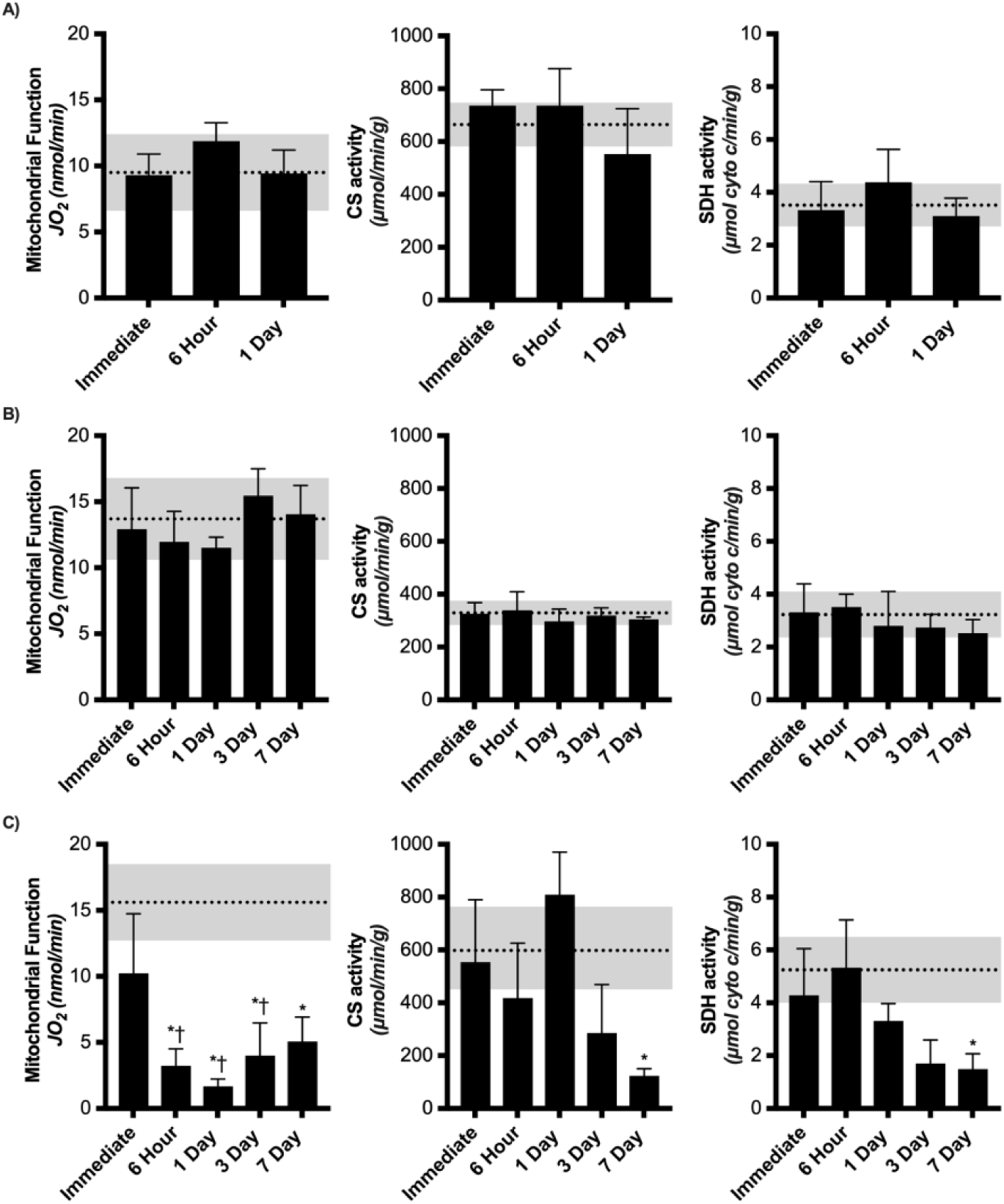
Mitochondrial function and content changes over time after different muscle stressors. Mitochondrial function was assessed by oxygen consumption measurements of permeabilized TA or EDL muscle fibers and mitochondrial content was assessed by mitochondrial enzyme assays of citrate synthase (CS) and succinate dehydrogenase (SDH) activity immediately, 6 hours, 1, 3, and 7 days after A) fatiguing protocol (n=12), B) Eccentric contraction-induced injury (n=20), C) and traumatic freeze injury (n=28). Dashed line represents average contralateral control limb and shaded grey regions are ± SD. Stressed limb data are presented as means ± SD. * Significantly different from uninjured, † significantly different from immediate injured.

**Figure 2.**
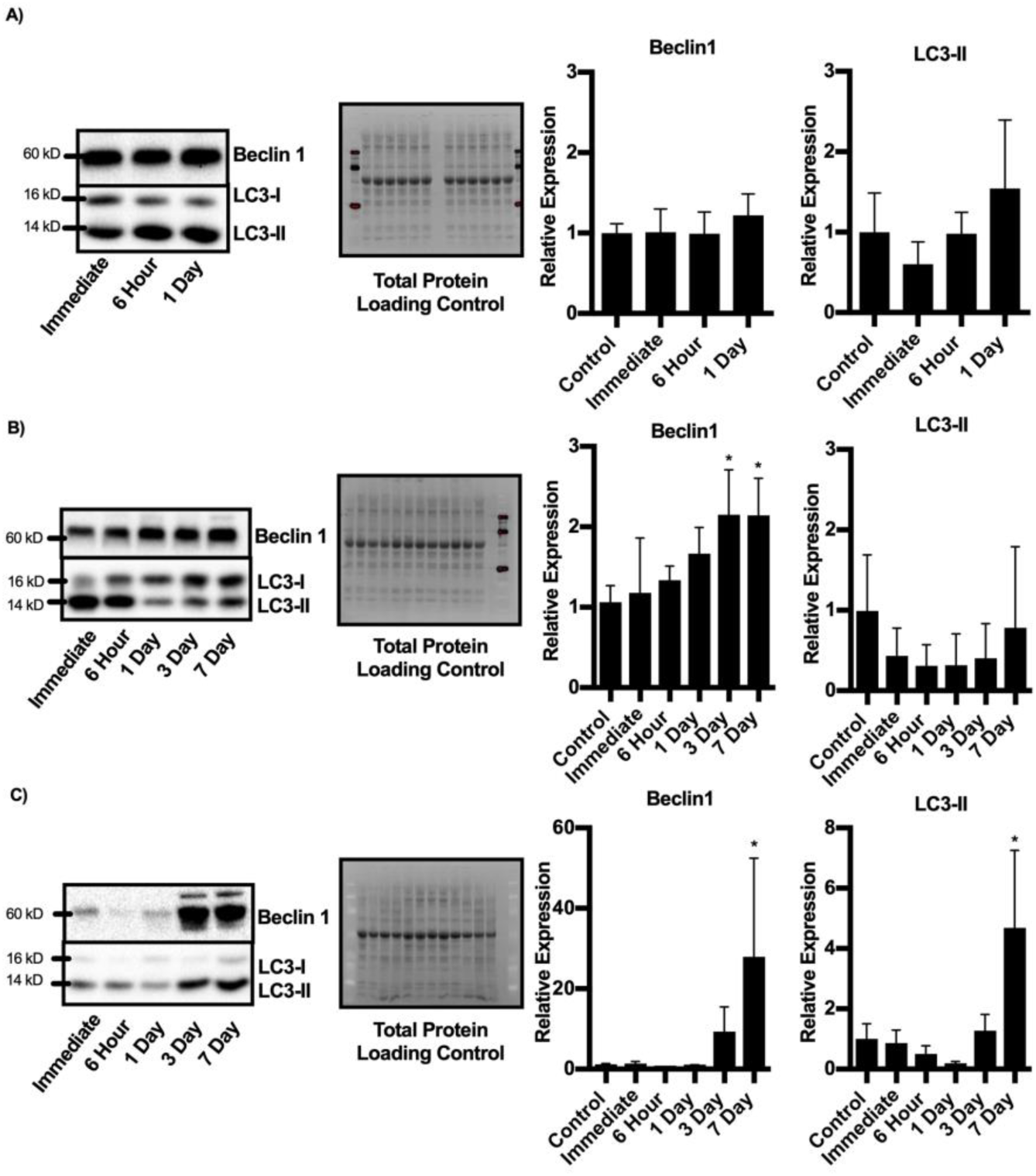
Autophagy related protein expression after different muscle stressors. Representative immunoblot and semi-quantitative analysis of Beclin1 and LC3II relative expression compared to control uninjured limb immediately, 6 hours, 1, 3, and 7 days after A) fatiguing protocol (n=12), B) Eccentric contraction-induced injury (n=20), C) traumatic freeze injury (n=28). * significantly different from control limb. Blots are normalized to total protein as a loading control and presented as relative expression to control uninjured limbs. Data are presented as means ± SD.

The second cohort of wildtype C57BL/6J mice were used to analyze autophagy flux and mitochondrial-specific autophagy following traumatic freeze injury based on results from the first cohort. Unilateral freeze injuries were performed on all mice before randomization into two groups. One group (n=12) was used for an autophagy flux assay where half of the mice received chloroquine to inhibit lysosomal degradation (17) and the other half were treated with saline 7 days after injury. Injured and contralateral limbs were collected and immunoblots for autophagic flux (LC3 II accumulation) were performed. The second group (n=8) was sacrificed 7 days after injury and a differential centrifugation protocol was done on both injured and contralateral uninjured limbs to determine the accumulation of autophagy-related proteins in mitochondria-enriched versus cytosolic fractions.

The third cohort of mice included Ulk1 MKO and LM mice to test the necessity of Ulk1 for recovery of mitochondrial function after injury (n=20) (10, 16, 18). Prior to injury, peak-isometric dorsiflexion torque measurements were performed on Ulk1 MKO and LMs. Immediately following, mice underwent unilateral freeze injuries and peak-isometric dorsiflexion torque measurements were performed again 14 days post. Mice were sacrificed afterwards, and muscle tissue was harvested for mitochondrial function, mitochondrial content, and autophagy-related protein analyses. The selection of 14 days after injury was based on the results from cohort 1 showing mitochondrial function was less than one-third of uninjured at 7 days after injury (Fig. 1), and results from our previous studies indicating 14 days is sufficient to observe differences in mitochondrial content with insufficient autophagy (6, 27)

### Metabolic Fatiguing Protocol

Mice were anesthetized using 1-2% isoflurane in oxygen, and left hind limb was shaved and aseptically prepared. The foot was positioned into a foot-plate attached to the servomotor (Model 129 300C-LR; Aurora Scientific, Aurora, Ontario, Canada) where the ankle joint was adjusted to a 90° angle and secured at the knee joint. Platinum-Iridium (Pt-Ir) needle electrodes were inserted percutaneously on both sides of the peroneal nerve and the testing platform was maintained at 37°C throughout the optimization and muscle stressor protocols. Optimal muscle stimulation was achieved by finding peak-isometric torque of the ankle dorsiflexors (tibialis anterior (TA), extensor digitorum longus (EDL), extensor hallucis longus muscles) through increasing the current stimulating the peroneal nerve at a 200 Hz pulse frequency prior to executing the muscle stressor protocol. The fatiguing protocol consisted of 30 minutes of continuous 10Hz stimulation which was modeled after muscle activation during a 30 minute treadmill run (3).

### Eccentric Contraction-Induced Injury

Mice were prepared, optimal muscle stimulation was verified following the methods listed above, and eccentric contraction-induced injury protocol was executed as previously described (4, 5). Briefly, for the eccentric contraction-induced injury, the foot was passively moved from the 0° position (perpendicular to the tibia) to 20° of dorsiflexion. The ankle dorsiflexor muscles were stimulated at 200 Hz for a 100-ms isometric contraction followed by an additional 50-ms stimulation while moving from 20° dorsiflexion to 20° plantarflexion at an angular velocity of 2000°/s. Eccentric contractions were repeated every 10 seconds until a total of 100 electronically stimulated eccentric contractions were complete.

### Freeze Injury

Freeze injury was performed as previously described (39). Before surgery, mice were anesthetized using isoflurane and given a local anesthetic injection of buvipicaine (5mg/kg)(17). Afterwards, the left limb was aseptically prepared, a 1.5cm incision was made over the TA muscle, and a steel probe cooled with dry ice was applied to the belly of the TA for 10 seconds. Upon completion of the freeze injury, the incision was closed with nylon suture and mice were administered meloxicam (2mg/kg) for pain management immediately and again 12 hours after surgery (39).

### Oxygen Consumption Rates

Mitochondrial function was assessed in dissected permeabilized muscle fiber bundles from both the stressed and contralateral control limb using methods adapted from Kuznetsov et al. and as we have previously described (19, 34). To ensure we were testing homogenously stressed muscle fibers, entire EDL muscles were permeabilized for the eccentric contraction-induced injury and fatiguing protocol and TA muscle fibers were dissected from the affected area for the freeze injury group. Oxygen consumption rates were made through the use of a Clark-type electrode (Hansetech) kept at a constant 25°C with constant stirring. State III respiration was accomplished by addition of glutamate (10mM), malate (5mM), succinate (10mM), and ADP (5mM). Oxygen consumption rates during State III respiration were normalized to tissue mass loaded into chamber.

### Enzyme Assays

Both citrate synthase (CS) and succinate dehydrogenase (SDH) enzyme assays were performed to quantify mitochondrial content in the stressed and contralateral control limbs after muscle fatigue, eccentric contraction-induced injury, and freeze injury. The portion of muscle remaining after fiber dissection for oxygen consumption rates was weighed and homogenized in 33mM phosphate buffer (pH 7.4) at a muscle to buffer ratio of 1:40 using a glass tissue grinder. Citrate Synthase activity was measured from the reduction of DTNB overtime as previously described (27). Succinate Dehydrogenase activity was measured from the reduction of cytochrome c as previously described (12).

### Immunoblot

For autophagy-related protein content analysis, protein was extracted from stressed and contralateral control muscles. 25 μg of total protein was separated by SDS-PAGE, transferred onto a PVDF membrane, and immunoblotted as previously described (27). The following antibodies (Cell Signaling, Danvers, MA) were used: Ulk1 (1:10000), beclin-1 (1:1000), and LC3B (1:1000). Immunoblots were normalized to total protein in lane and quantified using Bio-Rad Laboratories Image Lab software (Hercules, CA) (8, 35, 44).

### Chloroquine Treatment

In order to measure autophagy flux after injury we used a lysosomal inhibitor, chloroquine, as recommended by the autophagy guidelines (17). Mice underwent freeze injuries as described above and recovered for 7 days. Two hours before sacrifice mice were given an intraperitoneal injection of chloroquine (65mg/kg) to inhibit autophagosome degradation. TA muscle tissue was harvested and immunoblots for LC3II quantification were carried out as described above.

### Differential Centrifugation

To obtain mitochondrial-enriched fractions and cytosolic fractions, differential centrifugation was performed on injured and contralateral uninjured TA muscles 14 days after injury as described (20). Briefly, muscles were homogenized in fractionation buffer [20 mM HEPES, 250 mM Sucrose, 0.1 mM EDTA, plus protease and phosphatase] in a glass tissue homogenizer at a 1:20 tissue to buffer ratio. Homogenates were then spun at 800×*g* for 10 min at 4°C, supernatant was removed and then spun at 9000×*g* for 10 min at 4°C. The supernatant was again removed and resuspended in an equal volume of 2x Laemmli buffer resulting in the cytosolic fraction. The remaining mitochondrial pellets were resuspended in fractionation buffer then spun at 11,000×*g* for 10 min at 4°C. Resulting mitochondrial-enriched pellets were resuspended in 20 μl of 2x Laemmli buffer resulting in the mitochondrial-enriched fraction. Both fractions were boiled for 5 min at 97°C, then frozen at −80°C until immunoblot analysis.

### Immunofluorescent staining for satellite cells

Satellite cell dynamics were evaluated as previously described (42). Briefly, injured muscles from Ulk1 MKO mice (n=3) and LM mice (n=3) were isolated at 10 days post-injury and subjected to cryo-sectioning. Muscle sections were stained with primary antibody, Pax7 (1:5; DSHB) and Ki67 (1:1000; Abcam) overnight at 4 degrees, followed with a secondary antibody stain at room temperature for 1 hour. To evaluate satellite cell dynamics, muscle cross-sections (at 3 representative levels) were used to enumerate the total number of satellite cells (Pax7+), proliferating satellite cells (Pax7+/Ki67+), and self-renewing satellite cells (Pax7+/Ki67-).

### Statistics

Differences in mitochondrial function, mitochondrial content and autophagy protein expression after different muscle stressors were analyzed by two-way repeated measures (RM) analysis of variance (ANOVA) with the repeated measures being the injured vs. uninjured contralateral control limb and the other factor being time. Autophagy flux immunoblots were analyzed by two-way RM ANOVA with the repeated measures being the injured vs. uninjured contralateral control limb and the other factor being treatment (saline or chloroquine). Differential centrifugation immunoblots were analyzed by two-way RM ANOVA with the repeated measures being the injured vs. uninjured contralateral control limb and the other factor being fraction (mitochondria-enriched or cytosolic). Ulk1 KO and LM comparisons were analyzed by two-way RM ANOVA with the repeated measures being the injured vs. uninjured contralateral control limb and the other factor being genotype. All data were required to pass normality (Shapiro-Wilk) and equal variance tests (Brown-Forsythe *F* test) before proceeding with the two-way RM ANOVA. Significant interactions were tested with Tukey’s *post hoc* test using JMP statistical software (SAS, Cary, NC) to find differences between groups. Group main effects are reported where significant interactions were not observed. An α level of 0.05 was used for all analyses and all values are means ± SD.

## Results

### Time course of mitochondrial dysfunction and content after different muscle stressors

Across all conditions of muscle stress, there was no effect of time on mitochondrial function or content in the contralateral control limb (p≥0.55) therefore only the collective mean (dashed line) and standard deviations (grey region) are represented in each panel of Fig. 1. There was no difference in mitochondrial function or enzyme content between stressed and contralateral control limbs at any time point after the metabolic fatigue protocol (Main Effect: Limb, p≥0.24, Fig. 1A). Similarly, there was no difference in mitochondrial function or content following the eccentric contraction-induced injury protocol (Main Effect: Limb, p≥0.17, Fig. 1B). Mitochondrial function was significantly decreased 6 hours after traumatic freeze injury (20% of uninjured), continued to decline to its lowest functional capacity one day after injury (12% of uninjured), and by seven days after injury had recovered to ∼34% of uninjured control limbs (Significant Interaction: p=0.006, Fig. 1C). Mitochondrial content was not significantly different from contralateral control limbs until day 7 post-injury (Significant Interaction, p≤0.021, Fig. 1C) suggesting a disproportionate loss of mitochondrial function early after freeze injury.

### Time course of autophagy induction after different muscle stressors

Immunoblots of Beclin1 and LC3II were analyzed to determine the time course of autophagy induction after different muscle stressors. There was no change in relative expression of Beclin1 or LC3II in the stressed limb compared to the control limb after the fatiguing protocol (p≥0.728, Fig. 2A). Beclin1 expression increased 2-fold at 3 days after the eccentric contraction-induced injury and remained elevated through 7 days post injury (Significant Interaction, p=0.014, Fig. 2B), however, no significant change was observed with LC3II (p=0.9023). In contrast, traumatic freeze injury resulted in a robust autophagy induction evident by a 28-fold increase in Beclin1 expression and a 5-fold increase in LC3II at 7 days after injury (p≤0.008, Fig. 2C).

### Autophagy flux response after traumatic freeze injury

Static measurements of autophagy-related proteins are a poor indicator of dynamic autophagy activity, therefore, we measured autophagy flux by quantifying LC3II accumulation after chloroquine (CQ) treatment in freeze injured muscle as this muscle stressor had the most robust autophagy response (Fig. 2) (17). LC3II expression was increased nearly 14-fold in the injured limb compared to the uninjured limbs at 7 days after injury as previously found (27) (Main effect: Injury p<0.0001, Fig. 3). Additionally, CQ treatment resulted in greater LC3II accumulation independent of injury (Main effect: Treatment p=0.0178, Fig. 3). These results suggest that autophagy flux increases with muscle injury but does not appear to coordinate with the robust response of autophagic machinery after traumatic freeze injury.

**Figure 3.**
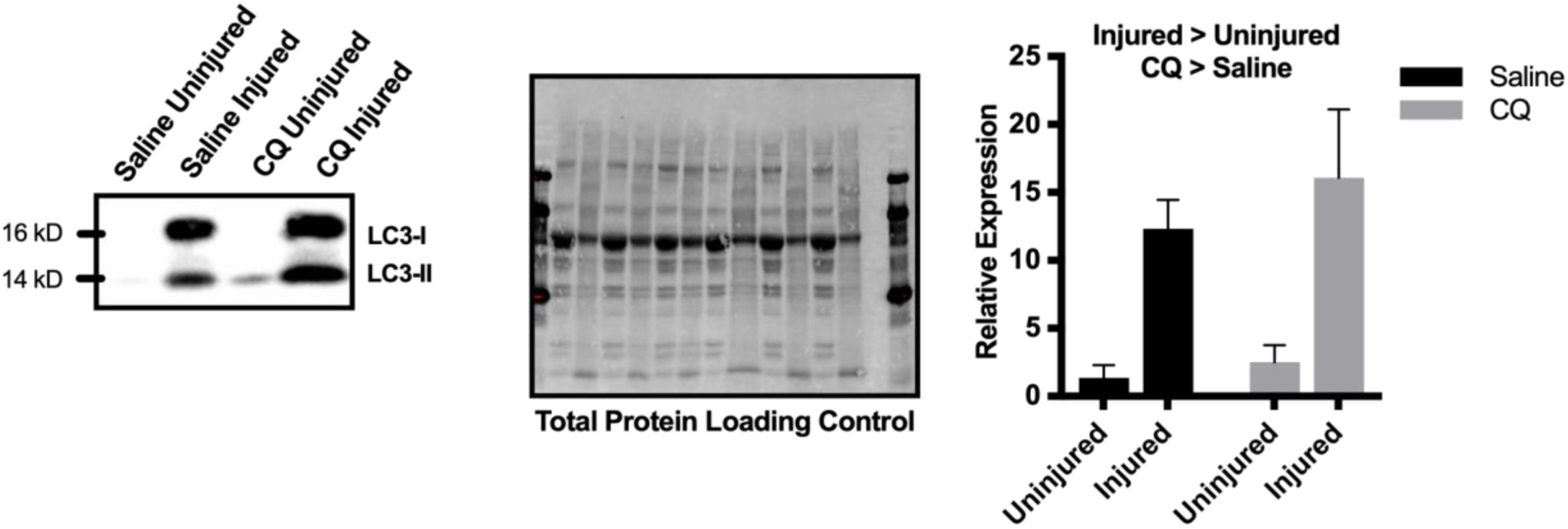
Autophagy flux after traumatic freeze injury. A) Representative immunoblot and semi-quantitative analysis of relative LC3II expression in both injured and contralateral limbs from mice treated with saline (n=6) or chloroquine (n=6) 7 days after freeze injury. Blots are normalized to total protein as a loading control and presented as relative expression to uninjured limbs from saline treated mice. Data are presented as means ± SD.

### Mitochondrial-specific autophagy after traumatic freeze injury

Because autophagy induction appeared to accompany mitochondrial dysfunction after freeze injury (Fig. 1 & 2), cytosolic fractions and mitochondrial-enriched fractions were subject to immunoblot analysis of LC3II to elucidate the extent of mitochondrial-specific autophagy. COXIV expression was increased 8-fold in the mitochondrial-enriched fractions compared to the cytosolic fractions (COXIV Main Effect: Fraction, p=0.049, Fig. 4). Interestingly, LC3II expression was 37 times greater in the injured mitochondrial-enriched fractions compared to the injured cytosolic fractions suggesting a large mitochondrial-specific autophagy response to traumatic freeze injury (Significant Interaction, p=0.017, Fig. 4), in agreement with previous reports of accumulation of autophagy-related proteins at the mitochondria after physiological muscle stress (6, 20).

**Figure 4.**
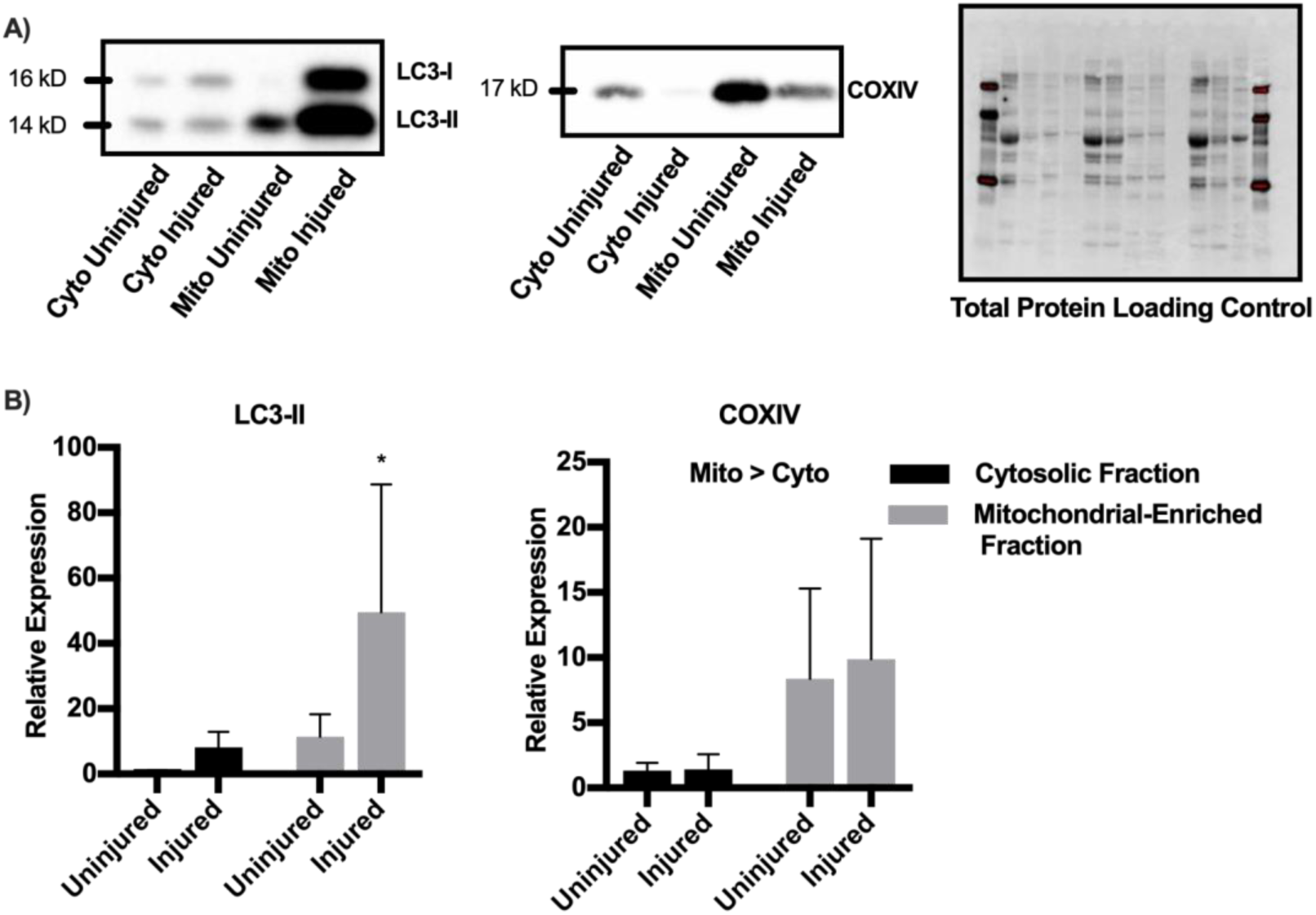
Autophagy induction in mitochondrial-enriched fractions after freeze injury. A) Representative immunoblots and B) semi-quantitative analysis of LC3II expression and COXIV expression in cytosolic and mitochondrial-enriched fractions from mice 7 days after freeze injury (n=8). Blots are normalized to total protein as a loading control and presented as relative expression to uninjured cytosol fraction. * significantly different from all other groups. Data are presented as means ± SD.

### Recovery of strength, mitochondrial function, and mitochondrial content in Ulk1 MKO mice after traumatic freeze injury

Mitochondrial-specific autophagy is mediated by the autophagy-related protein Ulk1 (6). We and others have investigated the role of Ulk1 following muscle stress and specifically mitochondrial stress (20, 27) however, whether Ulk1 is required for the recovery of mitochondrial function after injury has not been investigated. To ascertain the role of Ulk1-mediated autophagy in the recovery of mitochondrial function, we compared Ulk1 MKO and LM mice (6). Peak-isometric torque was significantly lower in Ulk1 MKO compared to LM mice independent of injury (−12%, Main Effect: Injury and Genotype, p≤0.012, Fig 5A). Mitochondrial function was decreased 43% in the injured limbs independent of genotype (Main Effect: Injury, p<0.0001, Fig 5B), and mitochondrial content was reduced by 34% and 58%, CS and SDH respectively, independent of genotype (Main Effect: Injury, p≤0.0006, Fig 5C, Fig 5D).

**Figure 5.**
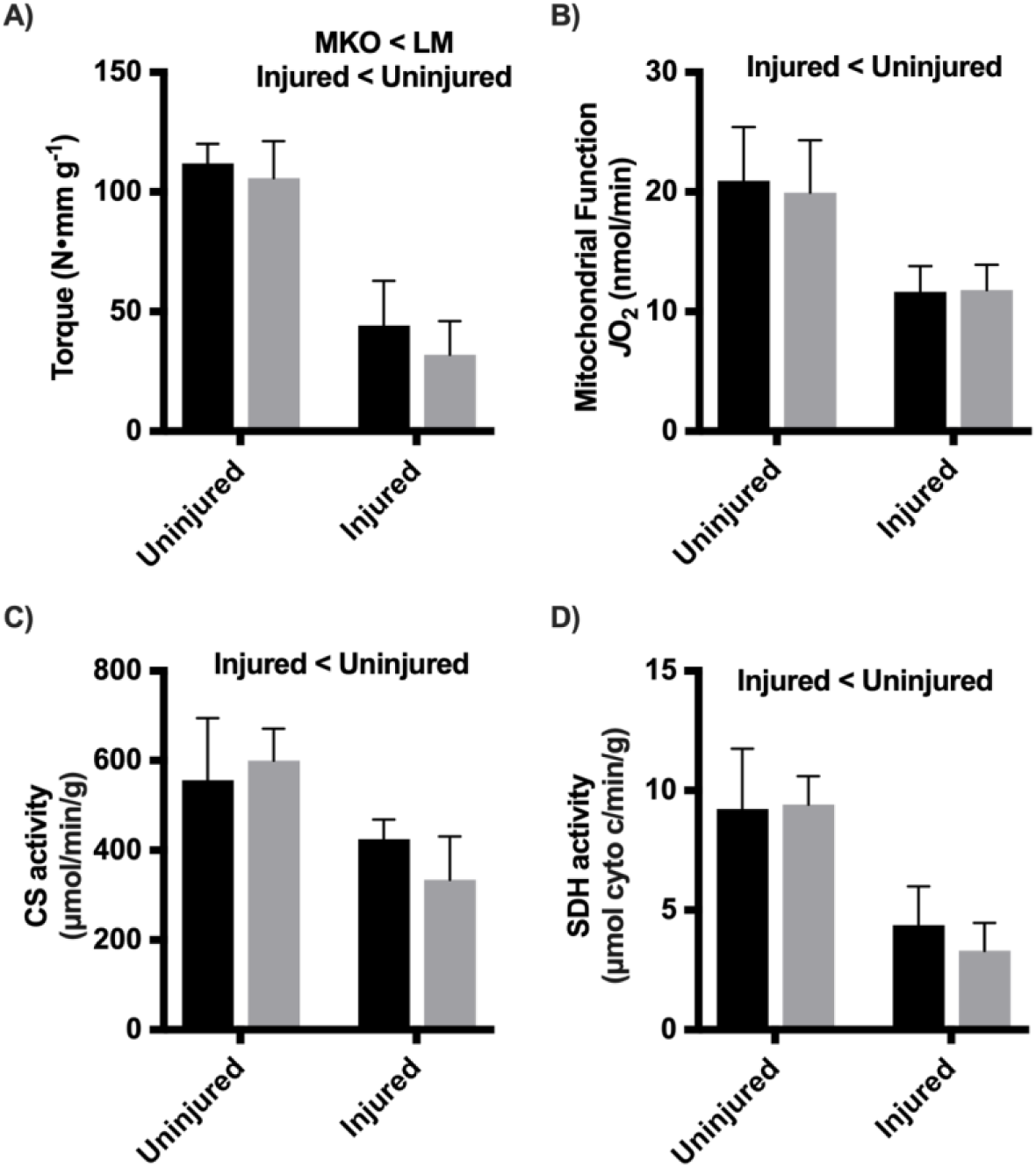
Muscle torque, mitochondrial function, and mitochondrial content before and after injury in MKO. A) Comparison of MKO (n=10) and LM (n=10) dorsiflexion muscle torque before (uninjured) and 14 days after freeze injury. B) Mitochondrial function assessed by oxygen consumption of permeabilized TA muscle in injured and contralateral uninjured limbs 14 days after freeze injury. C) Mitochondrial content assessed by Citrate Synthase activity and D) Succinate Dehydrogenase activity in TA muscles of LM and Ulk1 MKO mice in both injured and contralateral control limbs.

### Autophagy-related protein induction in Ulk1 MKO mice after freeze injury

LC3II expression increased more than 7-fold in the injured limbs compared to the uninjured limb, independent of genotype (Main Effect: Injury, p≤0.0001, Fig 6B). Additionally, Beclin1 expression increased more than 14-fold with injury, independent of genotype (Main Effect: Injury, p≤0.0001, Fig 6B). Prior to injury, Ulk1 MKO mice had no Ulk1 expression as expected, however after injury both the LM and Ulk1 MKO mice had similar levels of Ulk1 protein content (Main Effect: Injury and Genotype, p≤0.042, Fig 6B).

**Figure 6.**
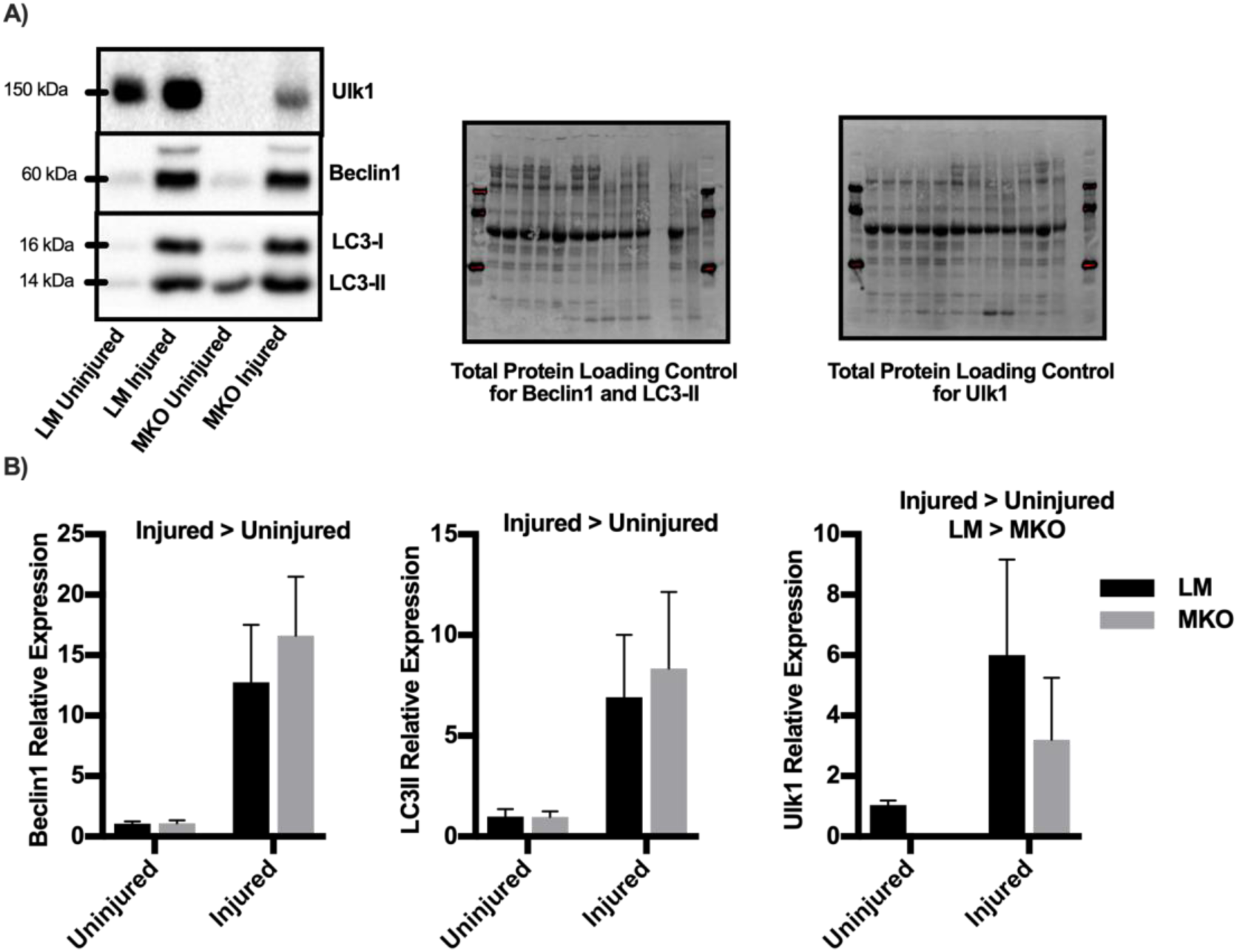
Autophagy related protein induction after traumatic freeze injury. A) Representative immunoblots and B) semi-quantitative analysis of Beclin1, LC3II, and Ulk1 protein expression in both injured and contralateral uninjured limbs of Ulk1 MKO (n=10) and LM (n=10) mice 14 days after freeze injury. Blots are normalized to total protein as a loading control and presented as relative expression to LM uninjured limbs. Data are presented as means ± SD.

### Impaired satellite cell proliferation in Ulk1 MKO mice

At the conclusion of our study we decided to explore satellite cell dynamics in Ulk1 MKO mice because: (i) the strength deficit in the Ulk1 MKO mice, which is in agreement with our previous reports (17), suggests Ulk1 knockout in myofibers impairs regenerative myogenesis and (ii) satellite cells are essential stem cells for regenerative myogenesis in skeletal muscle. 10 days post-injury there were a greater number of total and proliferating satellite cells in the freeze injured muscles of LM compared to Ulk1 MKO mice (p≤0.016, Fig. 7), and no difference between mice in the number of self-renewing satellite cells (p=0.140, Fig. 7).

**Figure 7.**
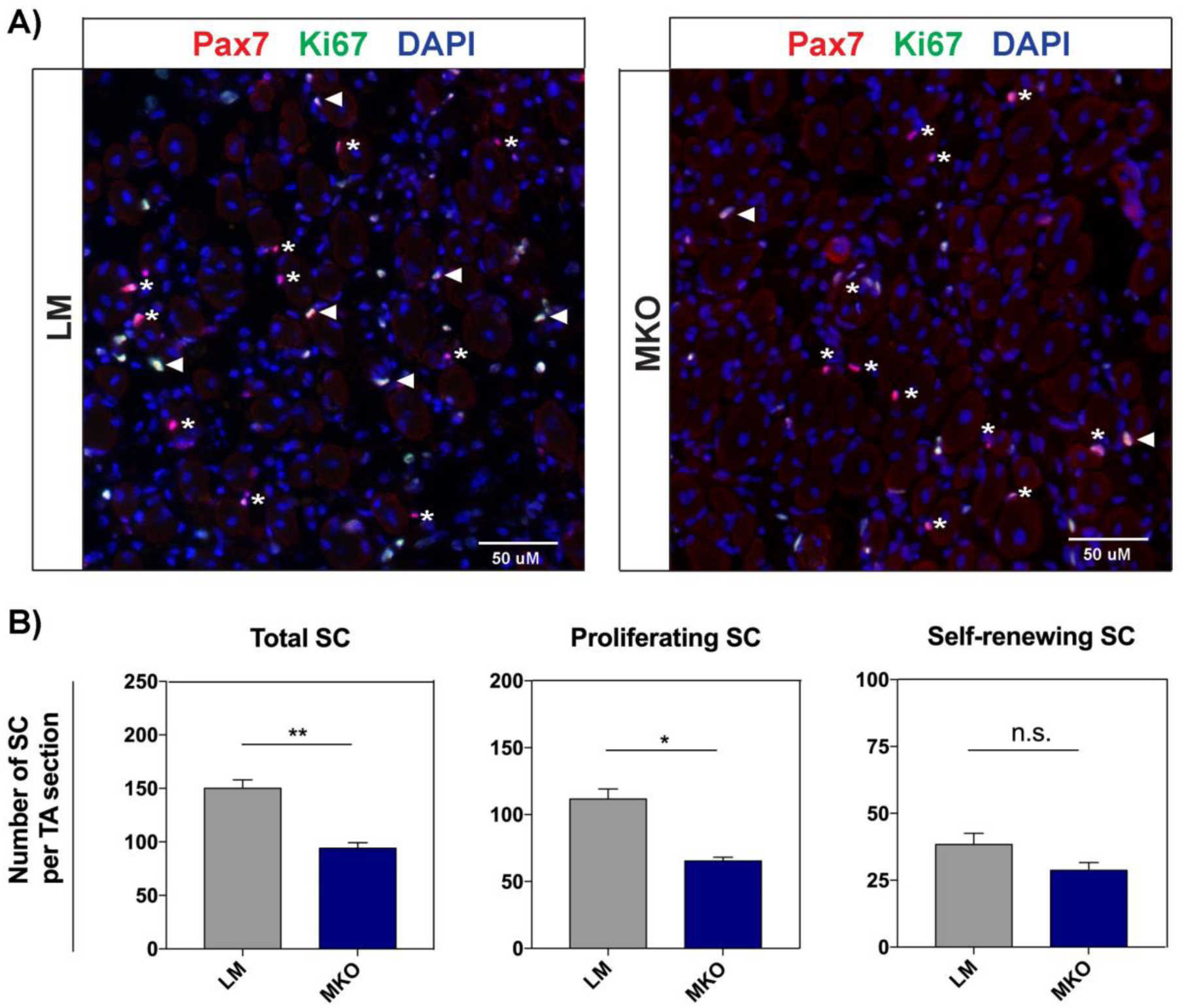
Muscle-specific Ulk1 knockout impairs satellite cell proliferation after traumatic freeze injury. A) Representative immunofluorescence imaging of muscle cross-sections from LM (n=3) and Ulk1 MKO (n=3) mice at 10 days after injury. Arrowheads: Pax7^+/Ki67+^ proliferative satellite cells. Asterisks: Pax7^+/Ki67-^ self-renewing satellite cells. B) Numbers of total, proliferating, and self-renewing satellite cells per muscle cross-section.

## Discussion

A primary goal for this study was to address several knowledge gaps in the field related to mitochondrial dysfunction after skeletal muscle stress, and the role of autophagy in mediating a response between the two. Mitochondria are appreciated as contributing to the regenerative potential, plasticity, and overall quality of skeletal muscle, and, therefore, investigating the muscle fiber-mitochondrial relationship may produce important targets for rehabilitation and disease prevention. However, there appear to be inconsistencies in the literature regarding what types of muscle stressors elicit mitochondrial dysfunction (20, 30, 31, 33), as well as the extent to which autophagy is necessary for the timely repair of mitochondrial dysfunction after muscle stress (6). Herein, we relied upon oxygen consumption as the marker of mitochondrial function, enzyme activities of succinate dehydrogenase and citrate synthase as markers of mitochondrial content, and a muscle-specific Ulk1 knockout mouse to test the necessity of autophagy for the recovery of mitochondrial function and content.

The first knowledge gap we explored was the extent to which three often utilized muscle stressors (fatigue, eccentric contraction-induced injury, freeze injury) that produce a loss in muscle contractility (21, 39) and an autophagic response (2, 6, 27) will cause mitochondrial dysfunction, i.e., a decline in oxygen consumption. Reduced mitochondrial oxygen consumption was only observed after traumatic freeze injury and was not decreased until 6 hours after injury (Fig. 1) in stark contrast to the immediate ∼70% loss in force production (39). Fatiguing exercises and eccentric contraction-induced injury have been reported to cause mitochondrial dysfunction (20, 26) in contrast to our findings. A likely explanation for these conflicting findings may be the tools utilized to investigate mitochondrial dysfunction and/or damage. Specifically, immunohistological techniques like confocal microscopy used by Laker & Drake et al (20) are useful for mechanistic investigations into localized mitochondrial events but do not necessarily reflect changes across the entire mitochondrial network; whereas oxygen consumption measurements test the functional capacity of the entire mitochondrial reticulum but do not capture more nuanced physiology such as fission and fusion events. It is irrefutable that functional, biochemical, and immunohistological approaches can provide valuable insight into mitochondrial physiology, yet when appropriate, future studies may consider the culminative advantage of combining more than one technique to avoid further inconsistencies in the literature.

Previous muscle damage research, including our own, has failed to investigate the dynamic properties of autophagy after skeletal muscle injury. Specifically, our previous work (6, 27) and this current work (Fig. 2) shows that the autophagy-related protein response scales to the magnitude of muscle damage after injury, but it remains unclear the extent to which autophagy flux matches the increases in autophagy machinery. Autophagy flux is described as the rate at which the entire process of autophagy occurs; meaning how quickly autophagosomes form around damaged content, fuse with lysosomes and subsequently degrade (17). In contrast, an increase in autophagy machinery is a static measurement and is not always connected to an increase in autophagy flux. Static measurements of LC3II protein content are often used as proxies for autophagy flux but can actually indicate (i) an inhibition in lysosomal fusion to the autophagosome, (ii) inhibition of autolysosome degradation, (iii) an increase in the amount of autophagosomes (without an increase in flux), or (iv) an increase in autophagy flux (17). Therefore, in order to asses changes in autophagy flux after injury we used the lysosomal inhibitor chloroquine and measured LC3II accumulation after blocking autolysosome degradation. We found that autophagy flux does increase a modest amount (∼1.5-fold increase) but does not scale to the large influx of autophagy machinery (∼6-fold increase) we observed after traumatic freeze injury (Fig. 3). This disproportionately small increase in flux may represent a limiting factor, or bottleneck, in the ability of autophagy to quickly clear away damaged proteins and organelles, and serve as a novel target to expedite the healing process in injured skeletal muscle.

We and others have previously reported that traumatic muscle injury results in a loss of mitochondrial content (6, 9, 27, 36), and herein we report that it also results in a loss of mitochondrial function (Fig. 1). A remaining question was the extent to which mitochondrial-specific autophagy, sometimes referred to as mitophagy, participates in the clearing of dysfunctional mitochondria after traumatic muscle injury. Laker & Drake et al. demonstrated that mitochondrial-specific autophagy was critical for clearing damaged, i.e, ROS-producing mitochondria after muscle fatigue (20); therefore, it is logical to hypothesize that mitochondrial-specific autophagy may occur after a more severe muscle stressor such as traumatic muscle injury in order to clear away the dysfunctional mitochondria. To address this knowledge gap, we analyzed autophagy-related proteins in mitochondrial-enriched fractions and cytosolic fractions. We found a robust accumulation of autophagy-related protein localized to the mitochondria after freeze injury suggesting that autophagy does participate in the clearance of dysfunctional mitochondrial after traumatic muscle injury (Fig. 4). This is the first study linking autophagy to mitochondrial dysfunction in injured skeletal muscle and understanding the process of clearing dysfunctional mitochondria after muscle injury may provide targets to facilitate muscle recovery.

In order to specifically test the extent to which mitochondrial-specific autophagy is important for muscle recovery after traumatic muscle injury we utilized a Ulk1 MKO mouse model. We and others have reported that Ulk1 may play an important role in both mitochondrial function and strength recovery after injury (6, 27). Our initial results suggested that Ulk1 is not required for the recovery of mitochondrial function after freeze injury (Fig. 5); however, there is a major consideration worth noting. Our Ulk1 MKO mouse model is a *myogenin-Cre* driven gene knockout, meaning Ulk1 is not expressed in adult muscle fibers, but is present in Pax7-expressing satellite cells. Quiescent satellite cells are activated upon injury, proliferate, differentiate, and ultimately provide new myonuclei for the regenerating fiber leading to Ulk1 expression in the regenerated muscle fiber. Additionally, following traumatic muscle injury there are many other cell types that migrate into the injured territory to aid in muscle regeneration. These include inflammatory cells such as neutrophils and macrophages, fibro-adipogenic precursor cells (FAPs), fibroblasts, and endothelial cells all of which potentially express Ulk1 sufficient for autophagy induction (41). This premise is supported by our immunoblots showing Ulk1 protein content within the injured limbs of Ulk1 MKO mice (Fig. 6). This is a clear physiological limitation of this study and limits our ability to determine the necessity of Ulk1 for mitochondrial remodeling after traumatic injury. To circumvent this problem for future experiments, we are exploring the use of *Pax7*^*CreER*^ mouse lines to effectively knockout Ulk1 in satellite cells and adult muscle fibers.

In this study, our finding that Ulk1 MKO impairs satellite cell proliferation raises an intriguing question – how does muscle fiber autophagy indirectly affect satellite cell dynamics? Satellite cells are essential stem cells for muscle regeneration and after traumatic injury (e.g., freeze injury), satellite cells exit quiescence and proliferate to form myoblasts (43). Myofiber-derived FGF2 and FGF6 are important mitogens for satellite cells during muscle regeneration (1, 7, 11, 15, 28). One possibility is that Ulk1-dependent autophagy may be pivotal for FGF2/6 expression and secretion in damaged myofibers. Alternatively, Ulk1-dependent autophagy may contribute to the degeneration of damaged myofibers by autophagy induced cell death, which would be critical for setting the stage for satellite cell proliferation via timely recruitments of macrophages and FAPs. No matter what mechanism is involved, the observations in this study suggest an indirect positive influence of autophagy in myofibers on satellite cell proliferation, which may be therapeutically targeted in the future for improving muscle regeneration.

In conclusion, this work advances the field in three substantive ways. First, physiological muscle stressors that cause a decrease in muscle contractility and potentially result in mitochondrial stress do not always elicit a decline in mitochondrial function, as assessed via oxygen consumption. Second, autophagy flux does not scale to the increase in total autophagy machinery that follows traumatic muscle injury, and this may represent a critical bottleneck to address with targeted therapies to enhance the recovery of muscle function. Third, autophagy appears to participate in the clearance of damaged mitochondria following traumatic injury in line with what has been reported following non-injurious muscle stressors (20). Therefore, future investigations into the role of autophagy following muscle stress associated with mitochondria should strongly consider an evaluation of mitochondrial function to complement a localized analysis of mitochondria stress and the use of a lysosomal inhibitor to determine autophagy flux. Unfortunately, we were unable to fully determine the necessity of Ulk1 for timely recovery of mitochondrial function due to the limitation of the mouse model; however, we are intrigued by the strength deficits in the mice and potential crosstalk between Ulk1 in the adult muscle fiber and satellite cells during muscle regeneration.

## Acknowledgements

Research reported in this publication was supported by the National Institute of Arthritis and Musculoskeletal and Skin Diseases of the National Institutes of Health under Award Number 1R01AR070178.

## References

1. Anderson JE, Mitchell CM, McGeachie JK, and Grounds MD. The time course of basic fibroblast growth factor expression in crush-injured skeletal muscles of SJL/J and BALB/c mice. Experimental cell research 216: 325–334, 1995.

2. Ato S, Makanae Y, Kido K, Sase K, Yoshii N, and Fujita S. The effect of different acute muscle contraction regimens on the expression of muscle proteolytic signaling proteins and genes. Physiological reports 5: 2017.

3. Baltgalvis KA, Call JA, Cochrane GD, Laker RC, Yan Z, and Lowe DA. Exercise training improves plantar flexor muscle function in mdx mice. Med Sci Sports Exerc 44: 1671–1679, 2012.

4. Call JA, Eckhoff MD, Baltgalvis KA, Warren GL, and Lowe DA. Adaptive strength gains in dystrophic muscle exposed to repeated bouts of eccentric contraction. J Appl Physiol (1985) 111: 1768–1777, 2011.

5. Call JA, Warren GL, Verma M, and Lowe DA. Acute failure of action potential conduction in mdx muscle reveals new mechanism of contraction-induced force loss. J Physiol 591: 3765–3776, 2013.

6. Call JA, Wilson RJ, Laker RC, Zhang M, Kundu M, and Yan Z. Ulk1-mediated autophagy plays an essential role in mitochondrial remodeling and functional regeneration of skeletal muscle. Am J Physiol Cell Physiol ajpcell.00348.02016, 2017.

7. Chakkalakal JV, Jones KM, Basson MA, and Brack AS. The aged niche disrupts muscle stem cell quiescence. Nature 490: 355–360, 2012.

8. Collins MA, An J, Peller D, and Bowser R. Total protein is an effective loading control for cerebrospinal fluid western blots. Journal of neuroscience methods 251: 72–82, 2015.

9. Duguez S, Feasson L, Denis C, and Freyssenet D. Mitochondrial biogenesis during skeletal muscle regeneration. American journal of physiology Endocrinology and metabolism 282: E802–809, 2002.

10. Egan DF, Shackelford DB, Mihaylova MM, Gelino S, Kohnz RA, Mair W, Vasquez DS, Joshi A, Gwinn DM, Taylor R, Asara JM, Fitzpatrick J, et al. Phosphorylation of ULK1 (hATG1) by AMP-activated protein kinase connects energy sensing to mitophagy. Science (New York, NY) 331: 456–461, 2011.

11. Floss T, Arnold HH, and Braun T. A role for FGF-6 in skeletal muscle regeneration. Genes & development 11: 2040–2051, 1997.

12. Foltz SJ, Luan J, Call JA, Patel A, Peissig KB, Fortunato MJ, and Beedle AM. Four-week rapamycin treatment improves muscular dystrophy in a fukutin-deficient mouse model of dystroglycanopathy. Skeletal muscle 6: 20, 2016.

13. Garcia-Prat L, Martinez-Vicente M, Perdiguero E, Ortet L, Rodriguez-Ubreva J, Rebollo E, Ruiz-Bonilla V, Gutarra S, Ballestar E, Serrano AL, Sandri M, and Munoz-Canoves P. Autophagy maintains stemness by preventing senescence. Nature 529: 37–42, 2016.

14. Hardy D, Besnard A, Latil M, Jouvion G, Briand D, Thepenier C, Pascal Q, Guguin A, Gayraud-Morel B, Cavaillon JM, Tajbakhsh S, Rocheteau P, et al. Comparative Study of Injury Models for Studying Muscle Regeneration in Mice. PloS one 11: e0147198, 2016.

15. Kastner S, Elias MC, Rivera AJ, and Yablonka-Reuveni Z. Gene expression patterns of the fibroblast growth factors and their receptors during myogenesis of rat satellite cells. The journal of histochemistry and cytochemistry : official journal of the Histochemistry Society 48: 1079–1096, 2000.

16. Kim J, Kundu M, Viollet B, and Guan KL. AMPK and mTOR regulate autophagy through direct phosphorylation of Ulk1. Nature cell biology 13: 132–141, 2011.

17. Klionsky DJ, Abdelmohsen K, Abe A, Abedin MJ, Abeliovich H, Acevedo Arozena A, Adachi H, Adams CM, Adams PD, Adeli K, Adhihetty PJ, Adler SG, et al. Guidelines for the use and interpretation of assays for monitoring autophagy (3rd edition). Autophagy 12: 1–222, 2016.

18. Kundu M, Lindsten T, Yang CY, Wu J, Zhao F, Zhang J, Selak MA, Ney PA, and Thompson CB. Ulk1 plays a critical role in the autophagic clearance of mitochondria and ribosomes during reticulocyte maturation. Blood 112: 1493–1502, 2008.

19. Kuznetsov AV, Veksler V, Gellerich FN, Saks V, Margreiter R, and Kunz WS. Analysis of mitochondrial function in situ in permeabilized muscle fibers, tissues and cells. Nature protocols 3: 965–976, 2008.

20. Laker RC, Drake JC, Wilson RJ, Lira VA, Lewellen BM, Ryall KA, Fisher CC, Zhang M, Saucerman JJ, Goodyear LJ, Kundu M, and Yan Z. Ampk phosphorylation of Ulk1 is required for targeting of mitochondria to lysosomes in exercise-induced mitophagy. Nature communications 8: 548, 2017.

21. Le G, Lowe DA, and Kyba M. Freeze Injury of the Tibialis Anterior Muscle. Methods Mol Biol 1460: 33–41, 2016.

22. Lovering RM, Roche JA, Bloch RJ, and De Deyne PG. Recovery of function in skeletal muscle following 2 different contraction-induced injuries. Archives of physical medicine and rehabilitation 88: 617–625, 2007.

23. Lynch GS, Hinkle RT, Chamberlain JS, Brooks SV, and Faulkner JA. Force and power output of fast and slow skeletal muscles from mdx mice 6-28 months old. J Physiol 535: 591–600, 2001.

24. Magalhaes J, Fraga M, Lumini-Oliveira J, Goncalves I, Costa M, Ferreira R, Oliveira PJ, and Ascensao A. Eccentric exercise transiently affects mice skeletal muscle mitochondrial function. Applied physiology, nutrition, and metabolism = Physiologie appliquee, nutrition et metabolisme 38: 401–409, 2013.

25. Marzetti E, Calvani R, Cesari M, Buford TW, Lorenzi M, Behnke BJ, and Leeuwenburgh C. Mitochondrial dysfunction and sarcopenia of aging: from signaling pathways to clinical trials. The international journal of biochemistry & cell biology 45: 2288–2301, 2013.

26. Molnar AM, Servais S, Guichardant M, Lagarde M, Macedo DV, Pereira-Da-Silva L, Sibille B, and Favier R. Mitochondrial H2O2 production is reduced with acute and chronic eccentric exercise in rat skeletal muscle. Antioxidants & redox signaling 8: 548–558, 2006.

27. Nichenko AS, Southern WM, Atuan M, Luan J, Peissig KB, Foltz SJ, Beedle AM, Warren GL, and Call JA. Mitochondrial maintenance via autophagy contributes to functional skeletal muscle regeneration and remodeling. Am J Physiol Cell Physiol 311: C190–200, 2016.

28. Olwin BB, and Hauschka SD. Identification of the fibroblast growth factor receptor of Swiss 3T3 cells and mouse skeletal muscle myoblasts. Biochemistry 25: 3487–3492, 1986.

29. Pauly M, Daussin F, Burelle Y, Li T, Godin R, Fauconnier J, Koechlin-Ramonatxo C, Hugon G, Lacampagne A, Coisy-Quivy M, Liang F, Hussain S, et al. AMPK activation stimulates autophagy and ameliorates muscular dystrophy in the mdx mouse diaphragm. The American journal of pathology 181: 583–592, 2012.

30. Rattray B, Caillaud C, Ruell PA, and Thompson MW. Heat exposure does not alter eccentric exercise-induced increases in mitochondrial calcium and respiratory dysfunction. European journal of applied physiology 111: 2813–2821, 2011.

31. Rattray B, Thompson M, Ruell P, and Caillaud C. Specific training improves skeletal muscle mitochondrial calcium homeostasis after eccentric exercise. European journal of applied physiology 113: 427–436, 2013.

32. Sandri M, Coletto L, Grumati P, and Bonaldo P. Misregulation of autophagy and protein degradation systems in myopathies and muscular dystrophies. Journal of cell science 126: 5325–5333, 2013.

33. Silva LA, Bom KF, Tromm CB, Rosa GL, Mariano I, Pozzi BG, Tuon T, Stresck EL, Souza CT, and Pinho RA. Effect of eccentric training on mitochondrial function and oxidative stress in the skeletal muscle of rats. Brazilian journal of medical and biological research = Revista brasileira de pesquisas medicas e biologicas 46: 14–20, 2013.

34. Southern WM, Nichenko AS, Shill DD, Spencer CC, Jenkins NT, McCully KK, and Call JA. Skeletal muscle metabolic adaptations to endurance exercise training are attainable in mice with simvastatin treatment. PloS one 12: e0172551, 2017.

35. Vigelso A, Dybboe R, Hansen CN, Dela F, Helge JW, and Guadalupe Grau A. Gapdh and beta-actin protein decreases with aging, making Stain-Free technology a superior loading control in Western blotting of human skeletal muscle. J Appl Physiol (1985) 118: 386–394, 2015.

36. Wagatsuma A, Kotake N, and Yamada S. Muscle regeneration occurs to coincide with mitochondrial biogenesis. Molecular and cellular biochemistry 349: 139–147, 2011.

37. Wang X, Pickrell AM, Rossi SG, Pinto M, Dillon LM, Hida A, Rotundo RL, and Moraes CT. Transient systemic mtDNA damage leads to muscle wasting by reducing the satellite cell pool. Human Molecular Genetics 22: 3976–3986, 2013.

38. Warren GL, Ingalls CP, Lowe DA, and Armstrong RB. Excitation-contraction uncoupling: major role in contraction-induced muscle injury. Exercise and sport sciences reviews 29: 82–87, 2001.

39. Warren GL, Summan M, Gao X, Chapman R, Hulderman T, and Simeonova PP. Mechanisms of skeletal muscle injury and repair revealed by gene expression studies in mouse models. J Physiol 582: 825–841, 2007.

40. Wilson RJ, Drake JC, Cui D, Ritger ML, Guan Y, Call JA, Zhang M, Leitner LM, Godecke A, and Yan Z. Voluntary running protects against neuromuscular dysfunction following hindlimb ischemia-reperfusion in mice. J Appl Physiol (1985) 126: 193–201, 2019.

41. Wosczyna MN, and Rando TA. A Muscle Stem Cell Support Group: Coordinated Cellular Responses in Muscle Regeneration. Developmental cell 46: 135–143, 2018.

42. Xie L, Yin A, Nichenko AS, Beedle AM, Call JA, and Yin H. Transient HIF2A inhibition promotes satellite cell proliferation and muscle regeneration. J Clin Invest 2018.

43. Yin H, Price F, and Rudnicki MA. Satellite cells and the muscle stem cell niche. Physiological reviews 93: 23–67, 2013.

44. Zeitler AF, Gerrer KH, Haas R, and Jimenez-Soto LF. Optimized semi-quantitative blot analysis in infection assays using the Stain-Free technology. Journal of microbiological methods 126: 38–41, 2016.

